# The Integrator complex regulates microRNA abundance through RISC loading

**DOI:** 10.1101/2021.09.21.461113

**Authors:** Nina Kirstein, Sadat Dokaneheifard, Pradeep Reddy Cingaram, Monica Guiselle Valencia, Felipe Beckedorff, Helena Gomes Dos Santos, Ezra Blumenthal, Mina Masoumeh Tayari, Ramin Shiekhattar

## Abstract

MicroRNA (miRNA) homeostasis is crucial for the post-transcriptional regulation of their target genes and miRNA dysregulation has been linked to multiple diseases, including cancer. The molecular mechanisms underlying miRNA biogenesis from processing of primary miRNA transcripts to formation of mature miRNA duplex are well understood^1-4^. Loading of miRNA duplex into members of the Argonaute (Ago) protein family, representing the core of the RNA-induced silencing complex (RISC), is pivotal to miRNA-mediated gene silencing^5-7^. The Integrator complex has been previously shown to be an important regulator of RNA maturation, RNA polymerase II pause-release, and premature transcriptional termination^8-11^. Here, we report that loss of Integrator results in global diminution of mature miRNAs. By incorporating 4-Thiouridine (s4U) in nascent transcripts, we traced miRNA fate from biogenesis to stabilization and identified Integrator to be essential for proper miRNA assembly into RISC. Enhanced UV crosslinking and immunoprecipitation (eCLIP) of Integrator confirms a robust association with mature miRNAs. Indeed, Integrator potentiates Ago2-mediated cleavage of target RNAs. These findings highlight an essential role for Integrator in miRNA abundance and RISC function.

---

MiRNAs are a class of non-coding RNAs of an average size of ∼22 nucleotides (nt), that are generated in two independent steps: i) primary miRNAs (pri-miRNAs) are processed by Microprocessor, composed of Drosha/DGCR8 to precursor (pre-miRNA) hairpins in the nucleus^2,3,12^, ii) after their export to the cytoplasm, Dicer/TRBP matures miRNA duplexes^13-15^. Further miRNA stabilization into RISC is critical for modulation of miRNA levels, and consequently the regulation of miRNA-target mRNA stability and translation. The Integrator complex is required for the cleavage of stalled RNA Polymerase II (RNAPII) transcripts, generating small promoter-associated RNAs^10^. The intriguing resemblance of a ∼20 nucleotide sub-population of this small RNA class and miRNAs led us to investigate the effect of Integrator depletion on miRNAs.

## Integrator depletion leads to miRNA loss

We depleted Integrator subunits (INTS, Extended Data Fig. 1a,b) and assessed the levels of mature miRNAs (n = 205; corresponding to the 200 most expressed miRNAs in shControl treated cells, extended for Drosha-independent miRNAs) using small RNA sequencing (smRNA-seq). Strikingly, we observed a global miRNA loss following INTS1, -3, -6, and -11 depletion, while INTS7 knock-down did not affect miRNA steady state (Fig. 1a,b). We found the strongest effect in absence of INTS6 and INTS11 (Fig. 1a-d, Extended Data Fig. 1c-g), with Drosha-independent miRNAs also being reduced (e.g.: 5’m^7^G-capped miR-320a-3p^16^ or mirtrons miR-877-5p and miR1226-3p^17^; Fig.1e,f, Extended Data Fig. 1h,i). We confirmed these findings by Taqman-qPCR miRNA detection in induced shINTS6 or shINTS11 HeLa cells and siINTS6, siINTS11, or siDrosha transfected HEK293T cells (Extended Data Fig. 1j,k). Importantly, we did not detect differential expression of pri-miRNAs, either by assessing nascent transcripts using PRO-seq after INTS11 knock-down (Extended Data Fig. 2a), or by total RNA-seq (Extended Data Fig. 2b,c). Concomitantly, while we detected some fluctuations in the expression of components of miRNA machinery (Extended Data Fig. 2d), and in the abundance of the major miRNA-related proteins (Extended Data Fig. 2e), no consistent alterations were identified that could explain miRNA loss. Additionally, we did not observe any changes either in pri-miRNA processing by Drosha/DGCR8 (Extended Data Fig. 2f-h), or in the average lengths of mature miRNAs (Extended Data Fig. 2i). Finally, miRNA loss was independent of Integrator’s endonucleolytic activity, as ectopic expression of wild-type or catalytic inactive E203Q-mutant INTS11 rescued the phenotype in shINTS11 cells (Extended Data Fig. 2j-l). These results indicate that while the abundance of mature miRNAs was controlled by the Integrator, loss of Integrator did not impact the processing of primary or precursor miRNA.

**Fig. 1.**
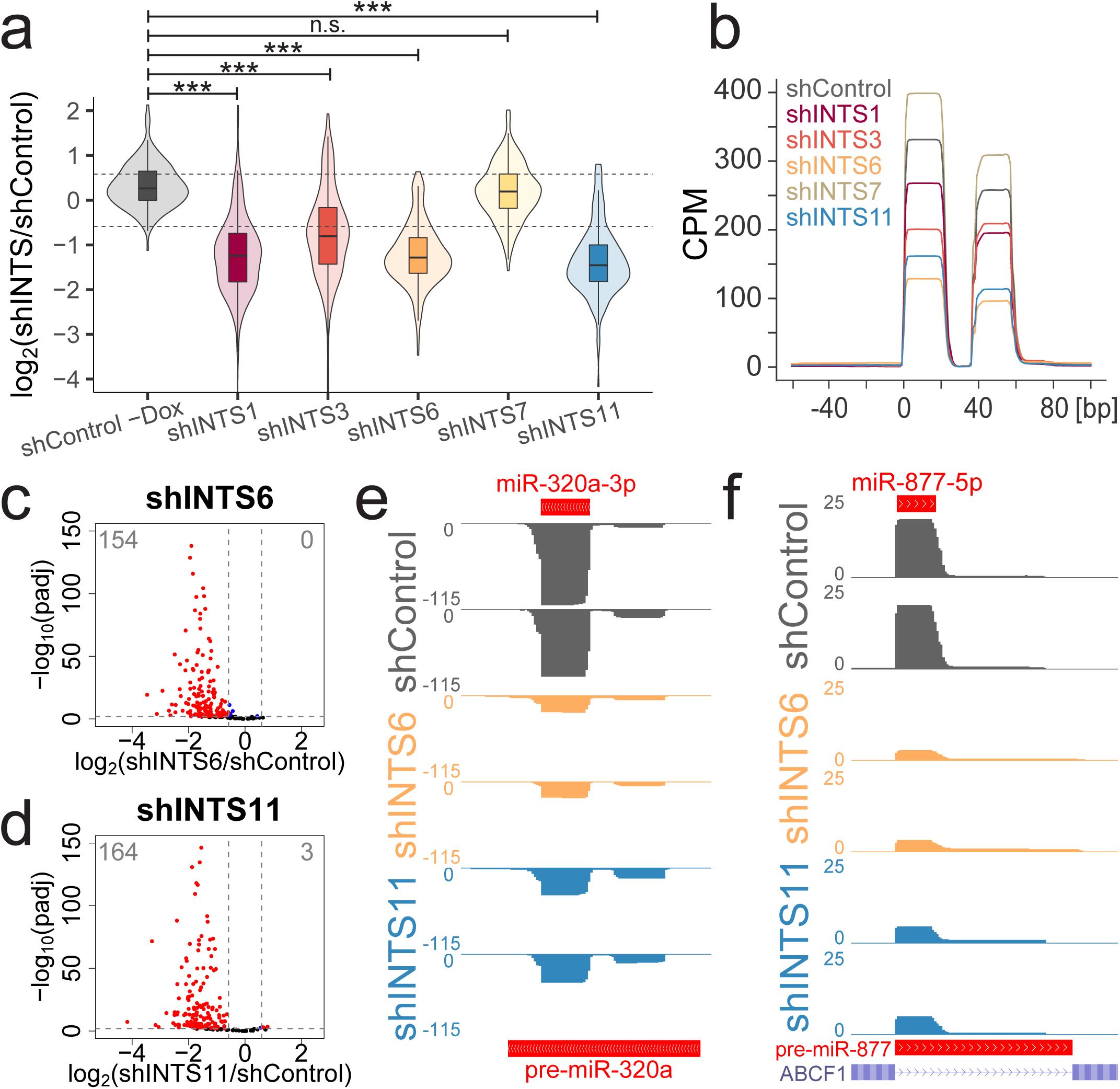
Integrator absence leads to global miRNA loss. **a**, Box- and violin plot depicting the log_2_ fold change of 205 expressed miRNAs determined by smRNA-seq in the indicated knock-down or uninduced shControl HeLa cells, calculated against induced shControl cells. *** p < 0.001, one-way ANOVA followed by Tukey’s post-hoc test. **b**, Global average smRNA-seq profiles around 112 5p-miRNAs aligned at their start site. **c**,**d**, Volcano plot comparing statistical significance and miRNA log_2_ fold change between control and knock-down cells. **c**, shINTS6. **d**, shINTS11. **e**,**f**, SmRNA-seq profiles at Drosha-independent miRNA loci. **e**, 5’-capped miR-320a locus. **f**, mirtron miR-877 locus.

## Integrator controls miRNA stabilization

To precisely pinpoint Integrator’s role in miRNA fate, we employed thiol (SH)-linked alkylation for metabolic sequencing of RNA (SLAM-seq) to determine smRNA dynamics^18^ following depletion of INTS6 or INTS11 (Fig. 2a). Briefly, 4-Thiouridine (s4U) was incorporated during transcription, which was subsequently carboxyamidomethylated (+Iodoacetamide, IAA), allowing to trace labeled miRNAs via their T>C conversion from a pool of labeled and unlabeled miRNAs (steady state). While 24h of s4U labeling did not affect miRNA levels (Extended Data Fig. 3a,b), global miRNA loss was still observed upon INTS6 or INTS11 depletion at steady state (Fig. 2b, Extended Data Fig. 3c,d). T>C labeled miRNAs were only detected in samples treated with s4U and IAA (Extended Data Fig. 3e-g), with a total number of 126 s4U labeled miRNA captured containing at least one T>C conversion (Extended Data Fig. 3h,i). Plotting the average RPM (reads per million) of T>C labeled miRNAs separated for guide (n=32) and passenger (n=32) strand allowed us to distinguish miRNA biogenesis (15min - 1h), initiation of RISC loading (1h - 3h), and miRNA stability reflected in differential abundance of guide and passenger strands seen between 3 to 24 hours in samples treated with shControl (Fig. 2c). Markedly, similar analysis following INTS6 (Fig. 2d) or INTS11 depletion (Fig. 2e) displayed the decreased separation of guide and passenger miRNA levels starting at 1h to 3h timepoints and extending throughout the 24 hours. This observation was confirmed when depicting the average of all detected miRNAs (n = 126; Fig. 2f, Extended Data Fig. 3j). We determined biogenesis (k_bio_) and accumulation rates (k_accu_) by linear regression of T>C labeled miRNAs on early (15min – 1h) or intermediate (1h - 6h) timepoints for either guide or passenger miRNAs (Fig. 2g). While knock-down of INTS6 appeared to result in an increase of passenger miRNA k_bio_, we did not detect any statistically significant change for either k_bio_ of guide or passenger miRNAs following INTS11 or INTS6 depletion (Fig. 2h, upper panel). As expected, k_accu_ was significantly larger for guide miRNAs as compared to that of passenger miRNAs in control knock-down condition, however depletion of INTS6 or INTS11 abrogated the difference between guide and passenger strand k_accu_ (Fig. 2h, lower panel). Similarly, median half-life estimations by single exponential saturation kinetics revealed reduced miRNA half-lives following shINTS6 treatment (t_1/2_=8.3h) compared to that of shControl (t_1/2_=12.4h), with greatest decrease observed following shINTS11 treatment (t_1/2_=1.2h), reflecting a defect in miRNA stabilization in absence of Integrator (Fig. 2i).

**Fig. 2.**
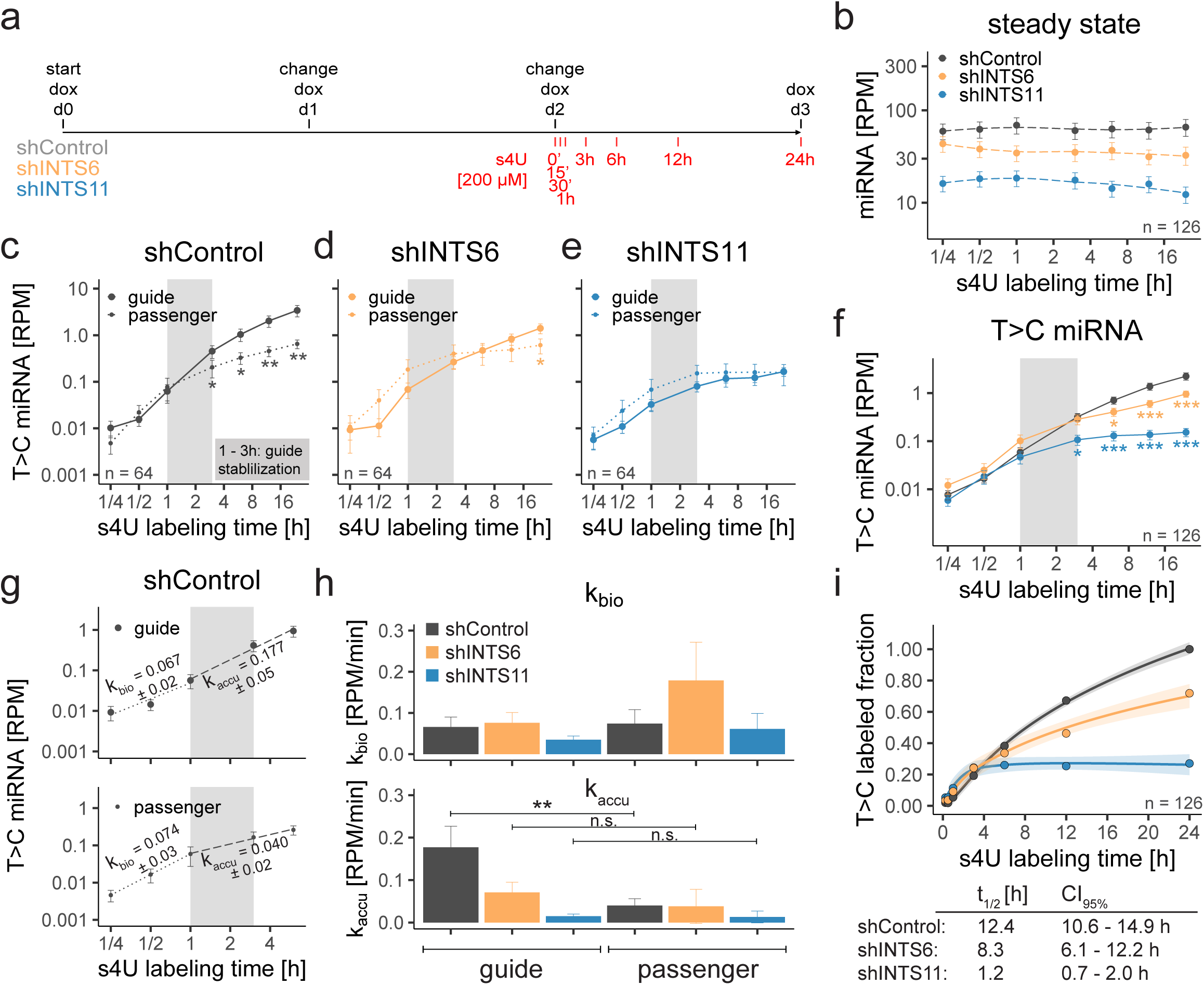
Depletion of Integrator abolishes miRNA stabilization. **a**, Scheme of INTS knock-down and s4U labeling. **b**, Steady state (unlabeled and T>C labeled) of miRNA expression over time. n = 126. **c-e**, T>C labeled miRNA abundance over time, separated for 32 guide or 32 passenger miRNAs. Mean ± SEM. * p < 0.05, ** p < 0.01, Mann-Whitney-Wilcoxon test. **c**, shControl. **d**, shINTS6. **e**, shINTS11. **f**, Combined T>C labeled miRNA abundance. **g**, Example of linear regression on shControl guide or passenger miRNAs. MiRNA biogenesis rates (k_bio_) determined from 15min to 1h, or accumulation rates (k_accu_) from 1h to 6h. Slope ± standard error is indicated. **h**, Histogram of k_bio_ and k_accu_. Mean ± SEM. ** p < 0.01, one-way ANOVA followed by Tukey’s post-hoc test on single miRNAs with k_bio_ or k_accu_ > 0. **i**, Single exponential saturation kinetics to calculate median half-life t_1/2_ [h] as depicted in table below including 95% confidence interval. Shades indicate SEM.

## Integrator loss abolishes RISC loading

MiRNA kinetics indicated that the absence of Integrator impaired miRNA stabilization, suggesting a role for Integrator in RISC loading. Indeed, analyses of miRNA associated with Ago2 using Taqman-qPCR following RNA-Immunoprecipitation (RIP), revealed the specific loss of Ago2-loaded miRNA after INTS11 or INTS6 depletion (Extended Data Fig. 4a,b). We next performed s4U labeling for 24h followed by Ago2 RIP to detect RISC-loaded miRNAs in shControl, shINTS6, and shINTS11 treated cells. We observed a decrease in steady state miRNA levels (Fig. 3a), confirming our previous RIP-qPCR results. Ago2 association of T>C labeled, newly generated miRNAs was significantly reduced upon INTS11 knock-down, with the same tendency for INTS6 (Fig. 3b, Extended Data Fig. 4c), reflecting an impairment of Ago2 loading. Importantly, miRNA loss was not rescued by concomitant overexpression of Ago2 (Fig. 3c, Extended Data Fig. 4d), further stressing the importance of Integrator during Ago2 loading.

**Fig. 3.**
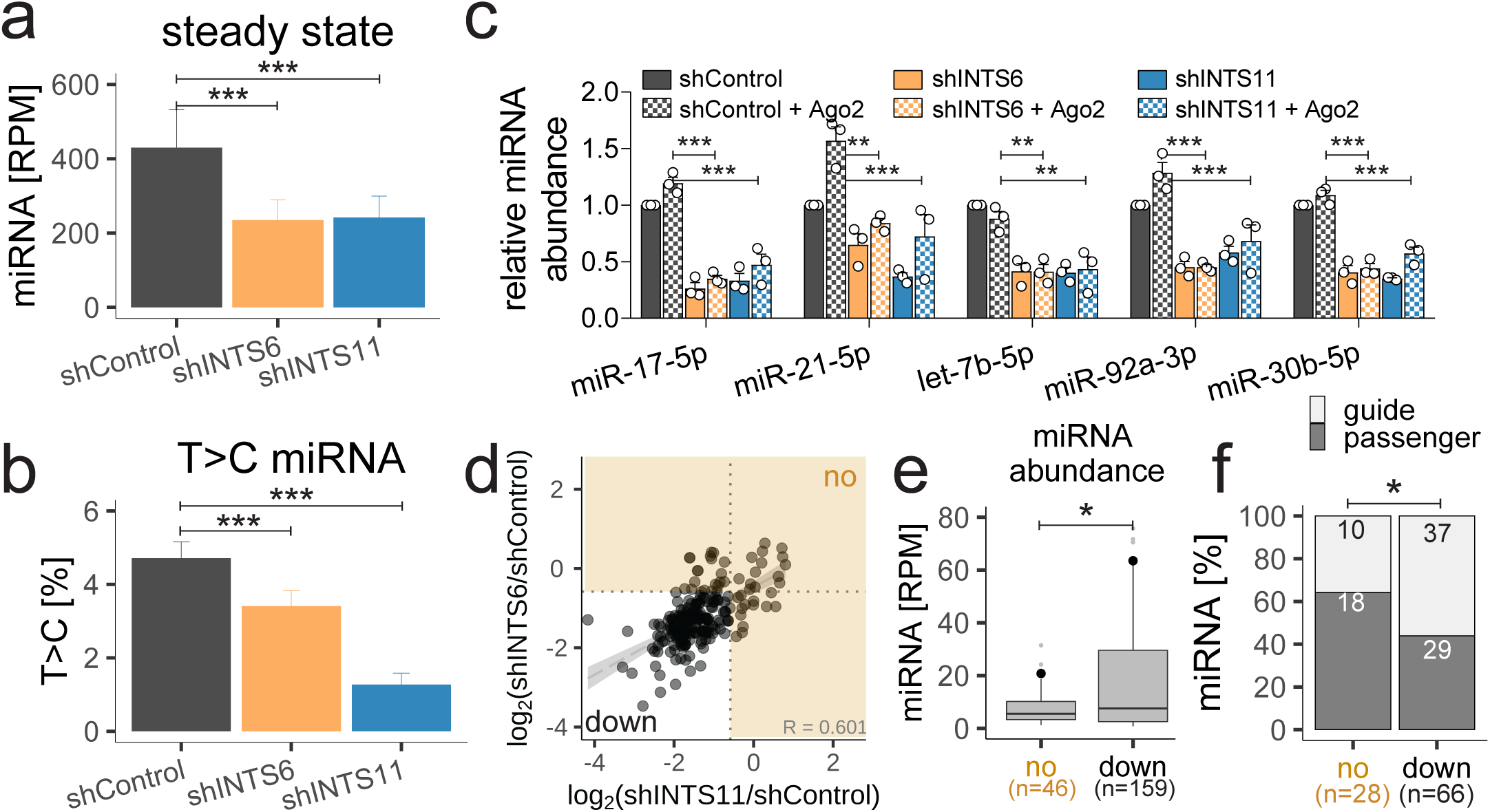
Integrator depletion abolishes Ago2 loading. **a, S**teady state miRNA abundance from Ago2 RIP after 24h + s4U (n = 122). **b**, T>C miRNA percentage. Mean + SEM. **c**, MiRNA TaqMan qPCR before and after Ago2 overexpression (see Extended Data Fig. 4d). MiRNA levels relative to ath-miR-159a spike-in and shControl. Mean + SEM, n = 3. ** p < 0.01, *** p < 0.001, one-way ANOVA followed by Dunnet’s multiple comparisons test. **d**, Scatter plot of the miRNA log_2_ fold change in shINTS6 and shINTS11 from Fig. 1c,d. Integrator-unregulated miRNAs are indicated by orange shaded area. (R: Spearman correlation coefficient). **e**, shControl miRNA abundance for unregulated (no) and down-regulated (down) miRNAs. * p < 0.05, Welch two sample t-test. **f**, Percentage of guide and passenger miRNAs. Absolute miRNA numbers are indicated. * p < 0.05, Fisher’s exact test.

## Loss of nuclear and cytoplasmic miRNAs

MiRNA stabilization in RISC is a cytoplasmic event^19-21^. While nuclear miRNA functions have previously been characterized^22^, they rely on active RISC transport to the nucleus^23,24^. Integrator has been described as nuclear complex with functions directly linked to active transcription^8-10,25-28^. Integrator is also present in the cytoplasm^29^. However, single INTS were two- (INTS11) to nine-fold (INTS6) enriched in the nucleus when examining equal amounts of HEK293T or HeLa nuclear or cytoplasmic extracts (Extended Data Fig. 4e). Furthermore, analyses of smRNA-seq from nuclear and cytoplasmic fractions following Integrator depletion revealed that miRNA loss was detected in both compartments, albeit stronger in the nucleus (Extended Data Fig. 4f). Consequently, Integrator acts on miRNA abundance in both cellular compartments.

Analysis of miRNA levels revealed a total of 46 miRNAs that were not significantly down-regulated following either INTS6 or INTS11 depletion (Fig. 3d, 17 miRNAs remain unregulated in absence of both INTS6 and INTS11). Interestingly, miRNAs down-regulated by Integrator depletion were significantly more abundant than miRNAs that were less affected by Integrator perturbations (Fig. 3e) and contained a higher proportion of guide miRNAs (Fig. 3f). These results further substantiate a cytoplasmic function of Integrator in miRNA stabilization into RISC.

## Integrator potentiates RISC function

A direct function of Integrator in Ago2 loading suggested an association of Integrator with mature miRNAs. We performed enhanced UV crosslinking and immunoprecipitation (eCLIP)^30^ optimized for the detection of miRNAs by increasing initial RNase concentrations, targeting INTS11 and its homolog CPSF73, a member of the cleavage and polyadenylation specificity factor (CPSF) complex. We detected miRNA binding by INTS11, but not CPSF73, confirming specific Integrator-miRNA interactions (Fig. 4a). We also detected INTS11 after anti-Flag affinity purification of Flag-Ago2, overexpressed in HEK293T cells (Extended Data Fig. 5a,b), supporting a transient interaction between Integrator and Ago2. Importantly, increasing concentrations of affinity-purified Integrator complex specifically enhanced recombinant Ago2’s ability to cleave a miRNA let-7a complementary sequence upon addition of duplex let-7a miRNA (Fig. 4b,c). Taken together, Integrator-dependent RISC loading not only modulates the abundance and stability of miRNAs but also functionally impacts gene silencing by RISC.

**Fig. 4.**
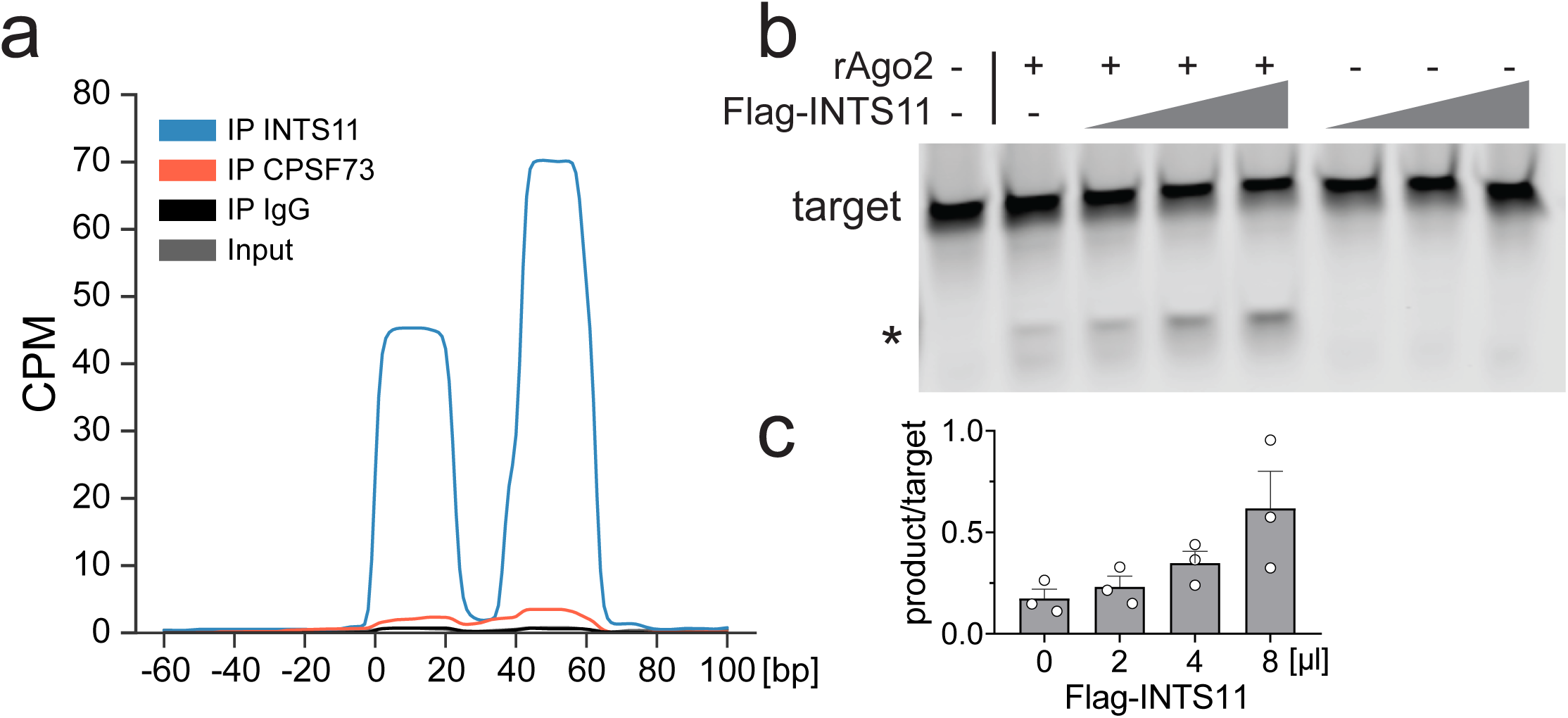
Integrator directly enhances Ago2 target cleavage efficiency. **a**, Global average of INTS11, CPSF73, IgG and size-matched input eCLIP profiles around 112 5p-miRNAs aligned at their start site. **b**, Ago2 cleavage assay in presence of dme-let-7a miRNA-duplex, 5’IRDye-700 labeled guide-complementary target RNA, Ago2 and increasing concentrations of affinity purified (Flag-INTS11) Integrator complex. * cleaved product. **c**, Quantification of the product/target ratio with increasing amounts of Integrator. Mean + SEM, n = 3.

## Discussion

Precise modulation of miRNA balance is crucial during cancer and development^31-33^. Integrator has previously been identified as critical for *Herpesvirus saimiri* pre-miRNA hairpin biogenesis^34,35^. While Integrator is enriched in the nucleus, we pinpoint a key function for Integrator complex in controlling human miRNA abundance in the cytoplasm, by directing their loading and consequently their stabilization in RISC. Given similar levels of miRNA reduction upon knock-down of INTS1, -3, -6, and -11, it is likely that multiple Integrator subunits endowed with RNA interaction domains are involved in RISC loading. Indeed, using eCLIP we showed a robust association of INTS11 and mature miRNAs. Significantly, functional reconstitution of miRNA-mediated targeted cleavage by Ago2 revealed a critical function for Integrator in proper loading of duplex RNA in RISC. These studies widen the scope of function of Integrator beyond transcriptional control and highlight a role for this complex in post-transcriptional regulation of gene expression by modulating miRNA stability and abundance.

## Methods

### Cell lines

HeLa (ATCC, #CCL-2) and HEK293T (ATCC, #CRL-3216) cells were maintained in high-glucose Dulbecco’s modified Eagle’s medium (DMEM, Gibco, #11965-084), supplemented with 10% fetal bovine serum (FBS, Atlas Biologicals, #F-0500-D) and the respective antibiotics, if required. Cells were regularly tested for mycoplasma. HeLa inducible shControl (containing an shRNA targeting GFP), shINTS1, and shINTS11 cells have been previously described in Gardini *et al*.^36^, shRNA-resistant N-terminal Flag-tagged WT and E203Q mutant INTS11 in Beckedorff *et al*.^10^. HeLa inducible shINTS3, shINTS6, and shINTS7 single clones were established by lentiviral infection with Tet-pLKO-puro vector (Addgene, #21915) containing the respective shRNA sequences (shINTS3: GCTGTGACCTCATTCGCTACA, shINTS6: ACCACTAATGATTCGATAATA, shINTS7: GCAGTAAAGAGACTTGCTATT) and 2.5 µg/ml puromycin (InvivoGen, #ant-pr) selection. shINTS stable cells were maintained in 2 µg/ml puromycin, WT and E203Q in presence of puromycin and 200 µg/ml G418 (InvivoGen, #ant-gn-5). shRNA expression was induced by adding Doxycycline (Selleckchem, #S4163) at 1 µg/ml for three days with medium replaced every 24 h.

### Cell transfections

HeLa cells were transfected with 20 nmol siControl (Ambion, #4390847), or siDrosha (Ambion, s26491) using Lipofectamin RNAiMAX (Invitrogen, #13778030) according manufacturer’s instructions. HEK293T cells were transfected with 30 nmol siControl, equimolar amounts of combined siINTS6 (s25483, s25484), siINTS11 (s29893, s29894, s29895), or siDrosha (s26490, s26491). For transient Ago2 overexpression in HEK293T cells, 3.2 µg of pFlag-CMV-Ago2 plasmid were transfected using Lipofectamine 2000 (Invitrogen, #11668019). Cells were harvested three days after transfection.

### Immunoblot detection

Whole-cell RIPA lysates were prepared in presence of Halt protease and phosphatase inhibitor cocktail (Thermo Scientific, #1861282) and proteins were separated by 4-15% Criterion TGX Stain-Free precast polyacrylamide gels (Biorad, # 5678084). After transfer on nitrocellulose membranes, we detected our protein of interest using the following antibodies: α-INTS1 (Bethyl Laboratories, #A300-361A), α-INTS3 (Sigma Prestige, HPA074391), α-INTS6 (Novus Biologicals, NB10086990), α-INTS7 (Bethyl Laboratories, A300-271A), α-INTS11 (Sigma Prestige, HPA029025), α-GAPDH (Abcam, ab8245), α-Drosha (Abcam, ab12286), α-Dicer (Abcam, ab14601), α-DGCR8 (Abcam, ab90579), α-Ago1 (Cell Signaling, D84G10), α-Ago2 (Abcam, ab57113), α-Ago3 (Sigma-Aldrich, SAB4200112), α-Ago4 (Cell Signaling, D10F10), α-Lamin B1 (Proteintech, #66095-1-1g).

### Subcellular fractionation

Nuclear and cytoplasmic fractions of HEK293T or HeLa cells were prepared as described in Bhatt *et al*.^37^. Briefly, cells were lysed in cytoplasmic lysis buffer (10 mM Tris-HCl, 15 mM NaCl, 0.15% NP-40) and layered on a sucrose cushion (10 mM Tris-HCl, 15 mM NaCl, 24% sucrose w/v), then centrifuged at 3.500g for 10 min at 4°C. Nuclear protein were extracted from the pellet using RIPA buffer and nuclear and cytoplasmic inserted for immunoblot as described above.

### RNA isolation

Total RNA was extracted using Trizol reagent (Thermo Fisher Scientific, #15596026) according to the manufacturer’s instructions. Nuclear and cytoplasmic fractions were prepared as described above, Trizol LS reagent (Thermo Fisher Scientific, #10296010) was used for extraction of cytoplasmic RNA. Genomic DNA was removed by Turbo DNase treatment (Invitrogen, #AM1907).

### MiRNA detection by Taqman qPCR

10 ng total RNA containing 5 pM ath-miR-159a spike-in was reverse-transcribed using Taqman Advanced miRNA cDNA synthesis kit (Applied Biosystems #A28007) and miRNAs were detected using specific probes for ath-miR159a (478411_mir), miR-17-5p (478447_mir), miR-21-5p (477975_mir), let-7b-5p (478576_mir), miR-320a-3p (478594_mir), miR-877-5p (478206_mir), miR-1226-3p (478640_mir), miR-92a-3p (477827_mir), miR-30b-5p, (478007_mir), miR-19a-5p (478750_mir), miR-182-5p (477935_mir), and RNU43_FAMMGB (Thermo Fisher, custom order) and SsoAdvanced Universal Probes Supermix (Biorad, #1725281). Relative miRNA expression was calculated against RNU43 or ath-miR-159a spike-in and shControl using ΔΔct method.

### Small RNA library preparation and genome mapping

Small RNA libraries were prepared using the SMARTer smRNA-seq kit (Takara, #635030) with 700ng total RNA. smRNA-seq from nuclear and cytoplasmic RNA was performed from 700 ng containing 35 ng (5%) of *Drosophila melanogaster* RNA as spike-in. Experiments were performed in two independent biological replicates that were sequenced together to avoid bias. For better comparability and to account for multiple sequencing runs, each including their own shControl, we merged the corresponding fastq files and randomly subsampled for 30 million reads before data processing. Sequencing reads were trimmed for adapter (AAAAAAAAAA) as recommend by SMARTer smRNA-seq kit (Takara, #635030) protocol using Cutadapt^38^ (v1.18) and reads shorter than 17 bp were omitted. Reads were aligned against human elements in RepBase (v23.08) with STAR^39^ (v2.5.3a) and the unmapped output then mapped against the human genome (hg19), allowing three mismatches and keeping all uniquely aligned reads. For UCSC Genome Browser visualization (https://genome.ucsc.edu/, ^40^), all tracks were normalized by CPM (counts per million) using deepTools2^41^ (v3.2.1).

### MiRNA detection and data analysis

Known mature miRNA were quantified using mirdeep2^42^ (v2.0.0.7) and the top 200 expressed miRNAs in shControl samples were selected and extended by Drosha-independent miRNAs. The final list of 205 miRNAs was analyzed in all data sets. Differential expression was calculated using DESeq2^43^ and R^44^ (version 3.6.1). Differentially expressed miRNA were determined by a cutoff of 1.5-fold and q-value of 0.01 (Supplementary Table 1). Significances were either calculated by DESeq2 or using one-way ANOVA testing followed by Tukey multiple pairwise comparisons in R. Graphics were generated using ggplot2^45^. Boxplots are represented with the median, the lower and upper hinges correspond to the first and third quartiles, the whiskers represent 1.5 × the inter-quartile range to both sides. Global average smRNA-seq profiles were based on bigwig files. MiRNA lengths were determined by mapping reads after removal of repetitive regions to miRNA precursors as described below (Small RNA SLAM-seq: Bioinformatic processing) and the mapped sequence lengths retrieved.

### Total RNA library preparation and genome mapping

Total RNA-seq libraries were generated using Truseq Stranded Total RNA library preparation kit (Illumina, #20020596) with 500 ng of DNase-treated RNA, including ribosomal RNA depletion. Sequencing was performed using Nova-seq to at least 50 million reads. Resulting fastq files were processed with Trimmomatic^46^ v0.32 and aligned to human genome (hg19) using STAR^39^ aligner v2.5.3a with default parameters. For UCSC Genome Browser^40^ visualization (https://genome.ucsc.edu/), all tracks were normalized by CPM (counts per million) using deepTools2^41^ (v3.2.1). RSEM^47^ v1.2.31 was used to obtain expected gene counts against the human Ensembl reference (release 87). Differential expression of shINTS compared to shControl was determined using DESeq2^43^. MiRNA-related genes were determined by selection relevant Gene Ontology (GO) terms^48^ containing “miRNA” (GO annotations: 0070883, 0070878, 0031054, 0035280, 1990428, 0035196, 2000634, 0031053, 0035281, 2000631) and extracting a list of 42 unique gene names.

### Pre-miRNA determination

The miRNA precursor file was obtained from miRbase^49^ v22, extracting entries annotated as “primary transcript” (corresponds to pre-miRNA) and lifting over to hg19 using CrossMap^50^. The resulting file was crossed with our list of 205 expressed miRNAs (BEDTools^51^ intersect, v2.29.0) to keep only relevant entries (n=176; GSE178127: 176_precursor_cleaned_hg19.bed).

### Primary miRNA determination and quantification

To assess expression of primary miRNAs, we used RNA-seq from siDrosha transfected HeLa cells as basis for new transcriptome assembly using StringTie^52^ (v2.0). We retrieved transcripts overlapping with annotated miRNA precursors of interest (see above) from the newly annotated transcript file, Ensembl annotation GRCh37.87 and GRCh38.99 (lifted to hg19 using CrossMap^50^). After curating the resulting annotation file manually, we obtained a final reference of expressed primary transcripts in HeLa cells (GSE178127: primir_final_annotation_v87_v99_StringTie_hg19_manualClean.gtf). We mapped shControl and shINTS RNA-seq data to the new primary reference and performed RSEM and DESeq2 as described above to assess differential pri-miRNA expression.

### Small RNA SLAM-seq: s4U treatment and carboxyamidomethylation

Small RNA SLAM-seq and data analysis was described in Reichholf *et al*.^18^. ShControl, shINTS6, and shINTS11 cells were seeded at a density of 1 × 10^6^ cells in 10 cm dishes at d-1 in DMEM medium (Gibco, #11965-084), supplemented with 10% FBS (Atlas Biologicals, #F-0500-D). shRNA induction was started at d0 by adding Doxycycline-containing medium (Selleckchem, #S4163, 1 µg/ml); the medium was changed every 24h. Two days after shRNA induction (corresponds to timepoint 0min), cells were additionally treated with 200 µM 4-Thiouridine (s4U, Cayman chemical, #16373). Parallel controls without s4U treatment were performed. Medium was exchanged every 3h during metabolic labeling to ensure homogenous incorporation and cells were kept from light exposure. At the respective timepoints (0min, 15min, 30min, 1h, 3h, 6h, 12h, 24h), cells were lysed directly on plate using TRIzol reagent (Thermo Fisher Scientific, #15596026) and samples were stored at -80°C until further processing. While protected from light, RNA was extracted according to the manufacturer’s instructions in presence of 0.1 mM DTT. Carboxyamidomethylation was performed as in described by Herzog *et al*.^53^. 40 ng RNA were treated with 10 mM Iodoacetamide (IAA, Sigma, #I1149-5g) dissolved in 100% Ethanol in presence of 50 mM NaPO_4_ and 50% DMSO at 50°C for 50 min. After quenching the reaction with 1 M DTT, RNA was Ethanol precipitated, followed by DNase treatment (Invitrogen, #AM1907). SmRNA libraries were prepared as described above using 800 ng of RNA containing 40 ng *Drosophila melanogaster* RNA as spike-in and sequenced with the Novaseq 6000 system (Illumina) to 40 – 90 million reads per sample.

### Small RNA SLAM-seq: Bioinformatic processing

SmRNA sequencing reads were treated as described above and mapped to repetitive regions. The resulting unmapped reads were mapped to a fasta file of 176 expressed miRNA precursors (as determined above) extended for 20 bp at their 3’end (GSE178127: 176_precursor_cleaned_hg19_ext20bp.fasta), while allowing for six mismatches using STAR v2.5.3a^39^. Minus strand miRNA precursors were annotated as reverse complement to allow all miRNAs to be treated as plus strand. Mapping smRNA-seq reads to the precursor file and applying a minimum threshold of 1 miRNA per million reads in all samples, we analyzed 126 miRNAs. Reads containing s4U-induced T>C conversions with a minimum base quality score of 27 were detected and analyzed as described in Reichholf *et al*.^18^ (https://github.com/breichholf/smRNAseq). Background (0min timepoint) was subtracted from the final RPM normalized reads. Median half-life was calculated based the T>C labeled fraction per timepoint, relative to shControl 24h, by nonlinear regression one phase decay analyses performed in GraphPad PRISM 8.0.

### Flag affinity purification

HEK293 stable cells overexpressing Flag-INTS11^8^ and Flag-Ago2^15^ were cultured in DMEM media (Gibco, #11965-084) containing puromycin and supplemented with 10% FBS (Atlas Biologicals, #F-0500-D). For Flag-INTS11 purification, nuclear lysate was extracted using buffer containing 20 mM Tris-HCl (pH 7.9), 1.5 mM MgCl2, 0.42 M NaCl, 25% glycerol, 0.5 mM DTT, 0.2 mM EDTA, 0.2 mM <u>PMSF.</u> For Flag-Ago2, cytoplasmic lysate was extracted using buffer containing 10 mM Tris-HCl (pH 7.9), 1.5 mM MgCl2, 10mM KCl, 0.5 mM DTT, 0.2 mM <u>PMSF.</u> Both complexes were purified from extract using anti-FLAG M2 affinity gel (Sigma). After washing twice with the buffer BC500 (20 mM Tris (pH 7.6), 0.2 mM ETDA, 10 mM 2-mercaptoethanol, 10% glycerol, 0.2 mM PMSF and 0.5 M KCl), and three times with buffer BC150 (20 mM Tris (pH 7.6), 0.2 mM ETDA, 10 mM 2-mercaptoethanol, 10% glycerol, 0.2 mM PMSF and 150 mM KCl), the affinity columns were eluted with FLAG peptide.

### RNA immunoprecipitation (RIP)

RIP was performed as described in Peritz *et al*.^54^. Protein A/G magnetic beads (Thermo Scientific, #26126) were rotated 16 hours with 10 µg Ago2 antibody (Abcam, ab57113). Cells were lysed in fresh polysome lysis buffer (100 mM KCl, 5 mM MgCl_2_, 10 mM HEPES (pH 7.0), 0.5% NP40, 1 mM DTT, 0.04 U/ml Superase-in RNase inhibitor (Ambion, #AM2694), Halt protease and phosphatase inhibitor cocktail (Thermo Scientific, #1861282)) and lysate was cleared by centrifugation. Protein concentration was determined using BCA protein assay and 350 µg protein lysate inserted in the IP reaction. After 16 hours, beads were directly lysed in 400 µl Trizol reagent (Thermo Fisher Scientific, #15596026) and RNA isolation was performed following the manufacturer’s instructions. For s4U-containing samples, IPs were performed in the absence of light and 2 pmol of synthetic ath-miR159a RNA was spiked-in after IP, to correct for isolation bias. RNA was isolated in presence of 0.1 mM DTT, resuspended in H2O, 1 mM DTT and the entire IP inserted in carboxyamidomethylation reaction as described above. After DNase treatment and final Phenol/Chloroform purification, equal volumes of IP RNA (400 – 600 ng RNA per IP) were subjected to smRNA library preparation as described previously. Sequencing data was treated as described above with the adjustment of mapping to ath-miR159a precursor prior to hsa-miRNA precursor mapping as described above, with the relative number of mapped reads serving as correction factor.

### eCLIP

eCLIP was performed in duplicates as previously described in Van Nostrand *et al*.^30^, optimized for the detection of mature miRNAs by increasing RNase I (Ambion, #AM2294) concentration from 40 U/ml to 200 U/ml. In brief, 2 × 10^7^ HeLa cells were crosslinked by UV-C irradiation (254 nm, 400 mJ/cm2) and lysed on ice followed by sonication. The lysate was subjected to RNase I (Ambion, #AM2294) digest (200 U/ml) in presence of murine RNase inhibitor (NEB, #M0314L) and 4 U/ml Turbo DNase (Ambion, AM2238). 4 μg of antibody (INTS11 (Sigma Prestige, #HPA029025), CPSF73 (Bethyl Laboratories, #A301-091A), rabbit IgG isotype control (Invitrogen, #02-6102)) was pre-incubated with Dynabeads M-280 Sheep Anti-Rabbit IgG (Invitrogen, #11204D) for 1 hour and added to the lysates for immunoprecipitation at 4°C for 16 hours. 2% of the lysate were removed and stored as size-matched input controls. Co-immunoprecipitated RNA was dephosphorylated, followed by on-bead 3’RNA adapter ligation using high concentration T4 RNA Ligase I (NEB, #M0204L). IP efficiency was verified by immunoblot of 20% of the IP samples. Input controls and 80% of the IPed protein-RNA complexes were run on a NuPAGE 4-12% Bis-Tris Plus Gels (Invitrogen, NP0321BOX), transferred to nitrocellulose membrane (Invitrogen, #IB23001) and the desired size range (protein size + 75 kDa) was cut from the membrane for IP and size-matched input samples. To extract RNA, nitrocellulose membranes were finely fragmented and treated with Urea/ Proteinase K, followed by acid phenol-chloroform extraction and purification using RNA Clean & Concentrator column cleanup (Zymo Research, #R1014). Input samples were also dephosphorylated and ligated to 3’RNA adapter. After reverse transcription (AffinityScript reverse transcriptase, Agilent, #600107), excess oligonucleotides were removed with exonuclease (ExoSAP-IT, Affymetrix, #78201), and the remaining RNA hydrolyzed by NaOH. A 3’ DNA Linker was ligated to the cDNA, and the resulting library was PCR amplified using Q5 Ultra II Master Mix (NEB, # M0544S). The library was size selected by agarose gel electrophoresis and column purified (Qiagen, MinElute, #28606). Single-end sequencing was performed to an average of 30 million reads per sample using Illumina NovaSeq 6000. Data was processed according to Van Nostrand *et al*.^30^ and https://github.com/YeoLab/eclip. After double adapter trimming (cutadapt^38^ v1.14), resulting reads were first mapped against the repetitive genome using STAR^39^ (v2.7.6a), the unmapped output was aligned against the human genome (hg19). PCR duplicates were removed by umi-tools^55^ (v1.0.0) and the samples were visualized in UCSC^40^.

### Ago2 cleavage assay

Ago2 cleavage assay was performed as previously described by Gregory *et al*.^6^ and the following modifications. Oligo RNAs were purchased from IDT: dme-let-7a-5p_guide (/5Phos/UGAGGUAGUAGGUUGUAUAGU), dme-let-7a-3p_passenger (/5Phos/UAUACAAUGUGCUAGCUUUCU), dme-let-7a_target (/5IRD700/UAUACAACCUACUACCUCAUU). MiRNA duplex was prepared by mixing equal volumes of both dme-let-7a-5p_guide and dme-let-7a-3p_passenger oligos in annealing buffer (10 mM Tris, pH 8, 50 mM NaCl, 1 mM EDTA), incubating at 95 °C for 3 min and cooling gradually to room temperature for 1 hour. Either 0.025 µg of recombinant Ago2 (rAgo2, Active Motif, #31486), or rAgo2 in combination with increasing concentrations of affinity-purified Flag-INTS11 were preincubated with the 5 nM of miRNA duplex in buffer containing 3.2 mM MgCl_2_, 1 mM ATP, 20 mM creatine phosphate, 0.2 U/μl RNasin, 20 mM Tris-HCl (pH 8), 0.1 M KCl, 10% glycerol for 30 min at 37°C. Then, 10 nM dme-let-7a_target was added and the cleavage reaction was incubated for 90 min at 37°C, stopped by adding Proteinase K for 30 min at room temperature. Samples were loaded onto a 15% TBE-Urea gels (Biorad, #4566055), visualized using Odyssey CLx Imaging System, and quantified by Image Studio Light (v5.2).

## Data availability

All sequencing data generated in this study is made available at the Gene Expression Omnibus (GEO). The accession number for the raw and processed data reported in this paper is GSE178127. Our previously reported PRO-seq data set is available under the accession GSE125535.

### Code availability

Code available upon request.

## Acknowledgements

We thank all members of the Shiekhattar lab for discussions. We thank Brian Reichholf for his comments on s4U small RNA SLAM-seq analysis. We thank the Sylvester Comprehensive Cancer Center Oncogenomics core facility for high-throughput sequencing. This work was supported by the University of Miami Miller School of Medicine, Sylvester Comprehensive Cancer Center and grants R01 GM078455 and DP1 CA228041 from the National Institute of Health to R.S. Research reported in this publication was funded by the National Cancer Institute of the National Institutes of Health under Award Number P30CA240139.

## Author contributions

N.K. performed small RNA-seq, small-RNA SLAM-seq, RIP-seq, and eCLIP (with the help of M.M.T.) experiments and bioinformatic analyses. S.D. performed Immunoblot, small RNA-seq, RNA-seq, RIP-qPCR, Ago2-rescue, Flag affinity purification, and *in vitro* Ago2 cleavage experiments. P.R.C. performed PRO-seq and *in vitro* experiments. M.V. established shINTS6 cell lines. F.B. and H.G. processed RNA-seq. E.B. set up small RNA-seq. N.K. and R.S designed the experiments and wrote the manuscript.

## Competing interests

The authors declare no competing interests.

## Additional information

### Supplementary Information

is available for this paper.

### Correspondence

should be addressed to R.S.

## Extended Data Figure Legends

**Extended Data Fig. 1.**
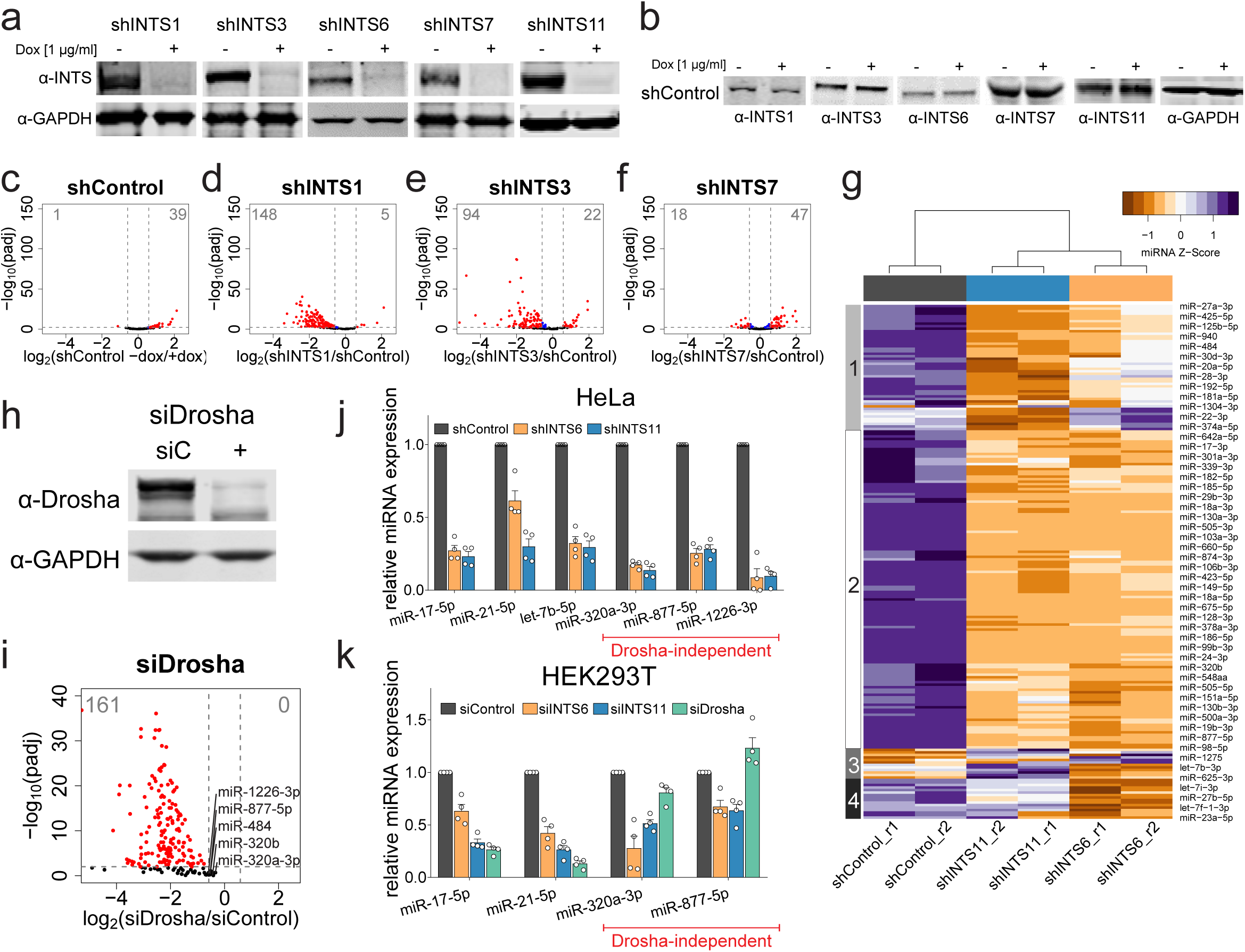
Drosha-independent miRNAs are down-regulated after INTS knock-down. **a**, Immunoblot detection of successful knock-down of INTS1, INTS3, INTS6, INTS7, and INTS11 before and after shRNA induction with Doxycyclin (Dox) at 1 µg/ml. GAPDH was used as loading control. **b**, Immunoblot of shControl before and after induction using the same INTS antibodies. **c-f**, Volcano plot comparing statistical significance and miRNA log_2_ fold change between control and knock-down cells. Significantly regulated miRNAs are depicted in red. **c**, uninduced shControl compared to induced shControl. **d**, shINTS1 compared to induced shControl. **e**, shINTS3. **f**, shINTS7. **g**, Heat map of normalized miRNA expression from shControl, shINTS6, and shINTS11 (Z-score of normalized read counts per row). Column and row orders were determined by unsupervised hierarchical clustering. **h**, Immunoblot detection of Drosha after siRNA knock-down in HeLa. **i**, Volcano plot comparing statistical significance and miRNA log_2_ fold change between siControl and siDrosha knock-down HeLa cells. Significantly regulated miRNAs are depicted in red. Drosha-independent miRNAs are indicated. **j**,**k**, Relative miRNA expression levels in **j**, HeLa shControl, shINTS6, and shINTS11 cells, or **k**, HEK293T cells transfected with siControl, siINTS6, siINTS11, or siDrosha. MiRNAs were detected by specific TaqMan probes for the indicated miRNAs and relative miRNA levels were calculated against RNU43 expression and shControl/siControl using ΔΔct method. Mean ± SEM, n = 4. Drosha-independent miRNA examples are indicated in red.

**Extended Data Fig. 2.**
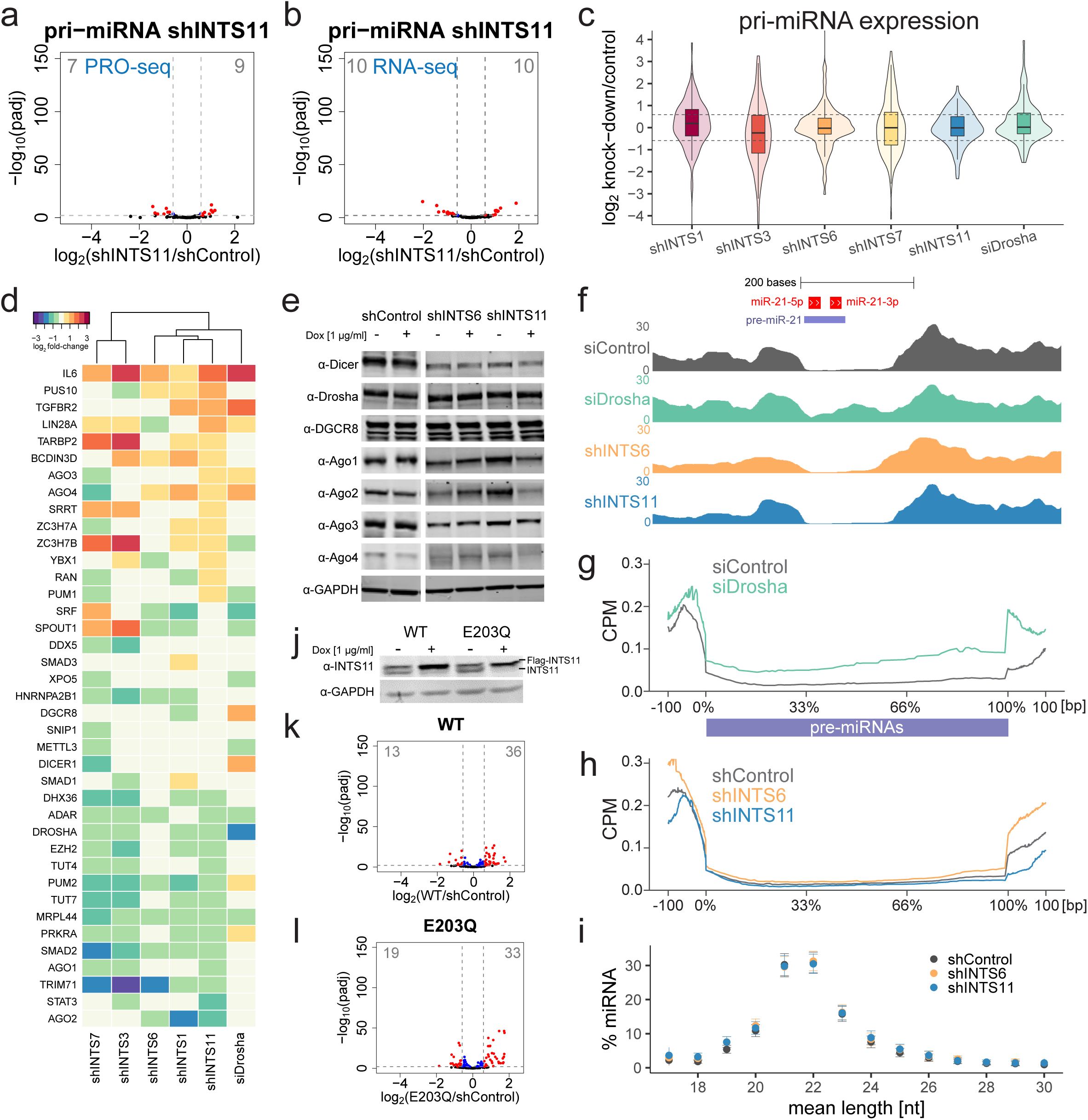
MiRNA loss does not depend on Integrator’s endonucleolytic cleavage activity. **a**,**b**, Volcano plot of statistical significance against log_2_ fold change between shControl and shINTS11 cells quantifying 109 pri-miRNAs in **a**, PRO-seq (transcriptional elongation) or **b**, total RNA-seq. Significantly regulated primary-miRNAs are depicted in red. **c**, Box- and violin plot depicting the log_2_ fold change of primary-miRNA expression obtained by total RNA-seq in the indicated knock-down cells, calculated against primary-miRNA levels in shControl or siControl cells. 109 pri-miRNAs were extracted from ENSEMBL or newly annotated transcripts based on siDrosha RNA-seq (see Material and Methods for details). **d**, Heat map of log_2_ fold changes in transcription levels of miRNA machinery-related genes after knock-down with the indicated sh/siRNAs calculated against sh/siControl. Gene names related to miRNA machinery were extracted from miRNA-containing Gene Ontology terms. Row order based on expression changes in shINTS11 cells, column order was determined by complete linkage hierarchical clustering. **e**, Immunoblot detection of the expression of miRNA biogenesis machinery and Argonaute proteins in shControl, shINTS6, and shINTS11 cells before and after induction. GAPDH was used as loading control. **f**, Example total RNA-seq profiles of indicated knock-downs depicting pre-miRNA excision at the miR-21 locus. Mature miR-21 are indicated in red, the annotated precursor is indicated in light blue. **g**,**h**, Cumulative total RNA-seq read densities across annotated pre-miRNAs ± 100 bp for **g**, siControl and siDrosha or h, shControl, shINTS6, and shINTS11 samples. **i**, Mean length [nt] percentage per miRNA ranging from 18 to 30 nucleotides detected in smRNA-seq of shControl, shINTS6, and shINTS11 cells. Mean ± SEM. **j**, INTS11 Immunoblot of shINTS11 cells stably expressing wild type INTS11 (WT) or catalytic mutant E203Q with and without shRNA induction. GAPDH was used as loading control. **k**,**l**, Volcano plot comparing statistical significance and miRNA log_2_ fold change detected by smRNA-seq between shControl and **k**, WT INTS11, or **l**, E203Q INTS11 cells in shINTS11 knock-down background. Significantly regulated miRNAs are depicted in red.

**Extended Data Fig. 3.**
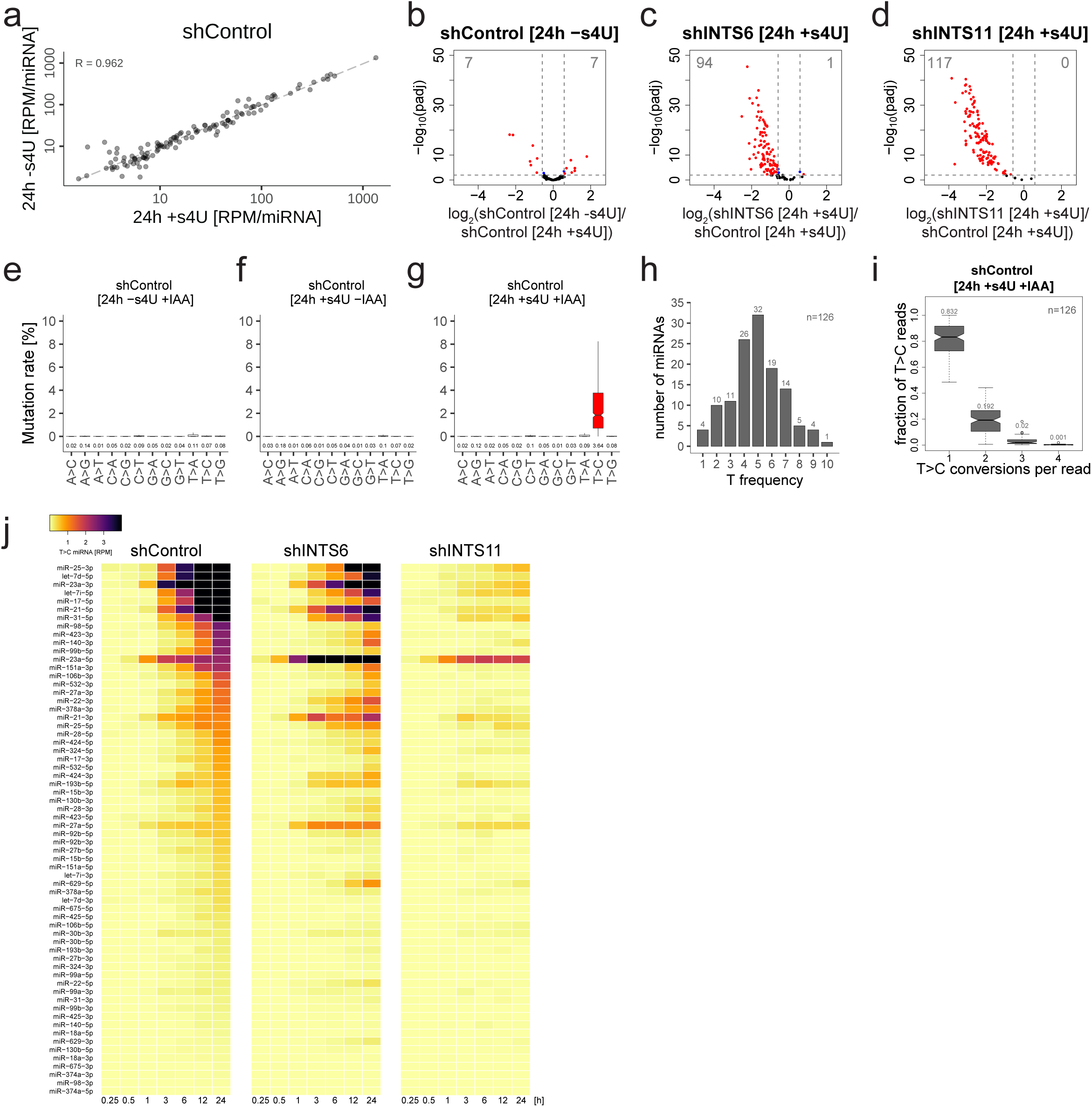
S4U labeling does not affect miRNA abundance. **a**, Scatter plot of steady state miRNA abundance [RPM] of 126 miRNAs in shControl samples with and without s4U treatment. Spearman correlation coefficient R is indicated. **b-d**, Volcano plot comparing statistical significance and miRNA log_2_ fold change between shControl cells [24h +s4U] and **b**, shControl cells control [24h -s4U]. **c**, shINTS6 cells [24h +s4U]. **d**, shINTS11 [24h +s4U]. Significantly regulated miRNAs are depicted in red, their numbers indicated on top. **e-g**, Conversion rates for every possible nucleotide conversion were detected for the miRNAs (positions 1-18 after background normalization) in shControl after 3d of Doxycycline treatment. **e**, Without s4U but with iodoacetamide (IAA) [24h –s4U +IAA]. **f**, [24h +s4U –IAA]. **g**, [24h +s4U +IAA]. Outliers were removed from representation; mean conversion rates are indicated below. **h**, Histogram representation of “T” frequency per miRNA in positions 1-18. n=126. **i**, Boxplot of the frequency of T>C conversion per read and per miRNA (n=126) in shControl cells after 24h s4U labeling and IAA treatment. The median fraction is indicated on top. **j**, Heatmap representation of the T>C miRNA expression [RPM] of 64 miRNAs (corresponding to 32 guide and passenger miRNA duplexes) during the time course of shControl (left panel), shINTS6 (middle panel), and shINTS11 (right panel).

**Extended Data Fig. 4.**
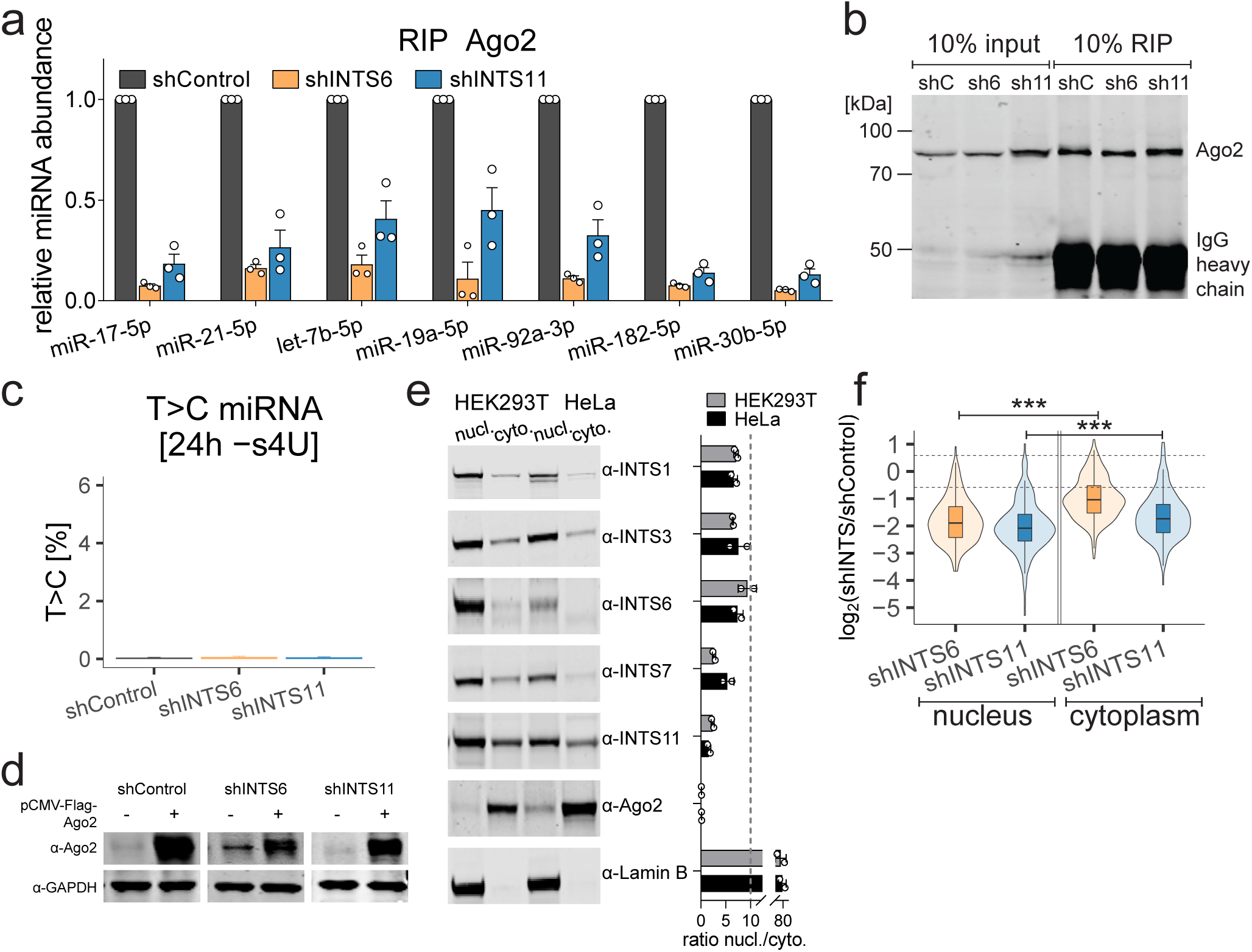
MiRNA loss is independent of subcellular localization. **a**, Ago2 RIP from shControl, shINTS6, and shINTS11 cells followed by Taqman-qPCR. MiRNA levels were normalized to shControl and ath-miR-159a spike-in. Mean ± SEM, n = 3. **b**, Immunoblot detecting Ago2 after example Ago2 RIP from induced shControl (shC), shINTS6 (sh6), and shINTS11 (sh11) cells. **c**, Percentage of T>C labeled miRNAs after Ago2 RIP from induced cells without s4U treatment. Mean ± SEM. **d**, Immunoblot detection of Ago2 in induced shControl, shINTS6, and shINTS11 cells, before and after transfection of pCMV-Flag-Ago2 plasmid. GAPDH servers as loading control. **e**, Left panel: Immunoblot detecting INTS as indicated from nuclear and cytoplasmic extracts from HEK-293T and HeLa cells. Lamin B serves as nuclear control. Right panel: Signal quantification and ratio of nuclear signal/ cytoplasmic signal of two independent experiments. **f**, Box- and violin plot depicting the log_2_ fold change of miRNA abundance per subcellular compartment obtained by smRNA-seq in the indicated knock-down cells, calculated against miRNA levels in induced shControl cells. 205 expressed miRNAs quantified by mirdeep2 were taken into account. Statistics were performed using one-way ANOVA followed by Tukey’s post-hoc test. *** p < 0.001.

**Extended Data Fig. 5.**
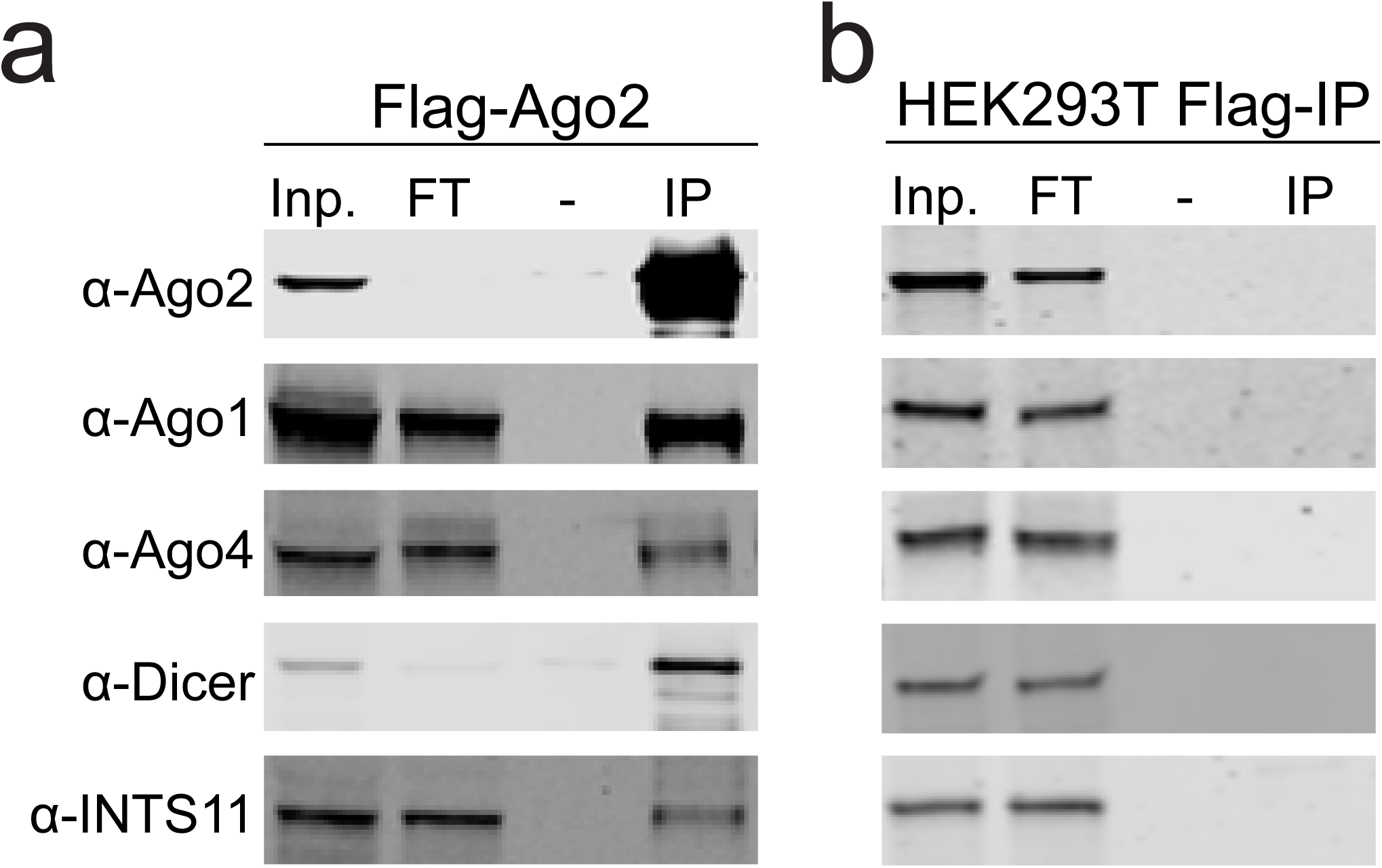
Integrator interacts with miRNA and Ago2. **a**, Flag affinity purification of Ago2 from HEK293T cells stably overexpressing pCMV-Flag-Ago2 probed for the indicated proteins. **b**, Control Flag affinity purification from parental HEK293T cells.

## Notes

### Competing Interest Statement

The authors have declared no competing interest.

